# Recurrent hyper-motif circuits in developmental programs

**DOI:** 10.1101/2024.11.20.624466

**Authors:** Miri Adler, Ruslan Medzhitov

## Abstract

During embryogenesis, homogenous groups of cells self-organize into stereotypic spatial and temporal patterns that make up tissues and organs. These emergent patterns are controlled by transcription factors and secreted signals that regulate cellular fate and behaviors through intracellular regulatory circuits and cell-cell communication circuits. However, the principles of these circuits and how their properties are combined to provide the spatio-temporal properties of tissues remain unclear. Here we develop a framework to explore building-block circuits of developmental programs. We use single-cell gene expression data across developmental stages of the human intestine to infer the key intra- and inter-cellular circuits that control developmental programs. We study how these circuits are joined into higher-level hyper-motif circuits and explore their emergent dynamical properties. This framework uncovers design principles of developmental programs and reveals the rules that allow cells to develop robust and diverse patterns.

## Introduction

Embryonic development is a well-orchestrated process in which cells divide, migrate, and differentiate into the diversified tissues that form the embryo. Starting from a single fertilized egg, cells use communication signals that they secrete and sense to regulate their cell fate decisions^1^. These signals, also known as morphogens, produce gradients and spatial patterns that inform the cells of their position and allow them to respond accordingly^2–6^. There are several families of morphogens that play key roles in shaping the spatial patterns in the embryo including the bone morphogenic proteins (BMPs)^7,8^, fibroblast growth factors (FGFs)^9^, Hedgehog (HH)^10^, WNT^11,12^ and Retinoic acid^13^. The production of these morphogens and their receptors is regulated by several transcription factor (TF) families that play critical roles in development, including HOX, GATA, ETS, SOX, PAX, TBOX, FOX, and E-proteins^14–22^. These morphogens and TFs regulate each other’s expression through intracellular regulatory circuits as well as cell-cell communication circuits, forming a complex network of interacting genes^23–29^ (Fig 1A). However, the rules that govern these interactions in regulating cell-fate decisions and forming tissue-level patterns remain poorly understood. Two complementary conceptual frameworks provide guiding principles governing cell fate decisions: Turing reaction-diffusion model and the concept of positional information^30–35^.

**Figure 1:**
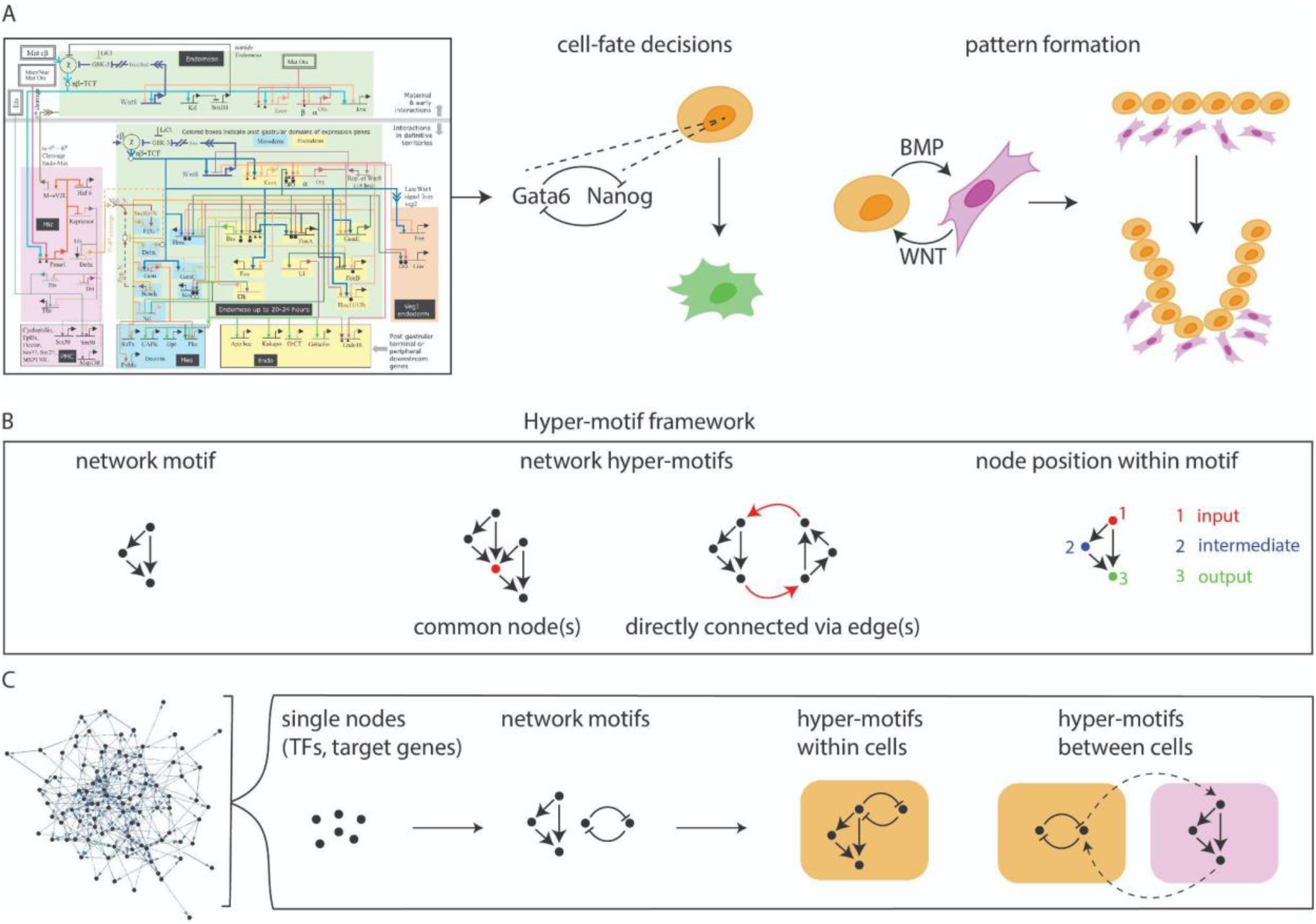
A network analysis framework to explore developmental programs. (A) Out of the multitude of regulatory interactions in gene regulatory networks (GRNs), what are the building block circuits of developmental programs for cell-fate decisions and pattern formation? (GRN adapted from Davidson et al., 2002^23^). (B) Definition of the hyper-motif framework: given two network motifs, their hyper-motifs are defined as combinations of motifs where the motifs share at least one node or interactions of motifs where the motifs are directly linked through at least one edge. Distinguishing different node positions within the motif is important to explore different hyper-motif structures. (C) A computational framework to analyze regulatory networks by identifying network motifs as building-block circuits and how they are joined together to form hyper-motifs within and between communicating cells.

Studies in model organisms have shed light on key pathways in developmental processes, where a detailed transcription network can be inferred in specific contexts from experimental data (as exemplified in Fig 1A). However, it is unclear which circuits out of the detailed transcription network are the ‘building blocks’ required for the proper development of the organism, and how their dynamical properties explain the wide range of observed tissue-level patterns^36^.

Previous theoretical and experimental work has defined network motifs, which are recurrent patterns, as the building blocks of biological networks^37,38^. Network motifs such as the feedforward loop (FFL) have been extensively studied and their dynamical properties were elucidated^39,40^. Seminal previous work has uncovered several families of network motif designs that can provide distinct behaviors in the interpretation of morphogen gradients^41^. How the integration of the individual network motifs within the complex regulatory network influences their functionality is not yet understood, however. Integration of small network motif patterns may generate emergent properties that cannot be explained by each motif alone. Understanding complex systems such as developmental regulatory networks therefore require the understanding of the rules of interaction between the systems’ building-block components. To address this, we recently defined hyper-motifs - higher-level network modules that describe how network motifs are embedded in large networks^42^. Given two network motifs, we consider two ways in which they can be directly joined to form a hyper-motif: having at least one node that is common to both motifs and being directly linked in the network through at least one edge (Fig 1B). Hyper-motifs can generate novel properties that are not observed in each motif alone and can also be used to silence certain motifs when not needed^43^. Such properties are critical to ensure accurate choice of cell fate, developmental timing and spatial patterning during embryogenesis. It is therefore important to understand how network motifs form these higher-level functional modules.

Here, we develop a network analysis framework to study the underlying principles of regulatory and cell-cell interactions that control developmental programs. Going beyond previous studies that focused on network motifs, we explore how the different network motifs in developmental programs are wired together within cells and between different cells through morphogen signaling pathways (Fig 1C). This framework can help explain how cellular interactions that are governed by a minimal set of regulators show diverse, complex and precise emergent properties.

## Results

### Enriched network motifs in the development of the human intestine

We analyze single-cell RNA-sequencing (scRNAseq) data from the human intestine across 9 time points, spanning 8-22 post-conceptual weeks (PCWs), from Fawkner-Corbett et al.^44^ (Fig 2A-B). We focus on developmental-related genes including developmental TFs, morphogens ligands, their antagonists, receptors and co-receptors. We used SCENIC^45^ to infer regulatory interactions between the developmental transcription factors and their target genes (Fig 2C, Methods). The networks we inferred are such where we sum over all regulatory interactions between genes that are expressed across different cell types. The size of the networks remains roughly constant across time, with the number of genes ranging between 220 and 275 and number of edges between 752-1045 (Fig S1).

**Figure 2:**
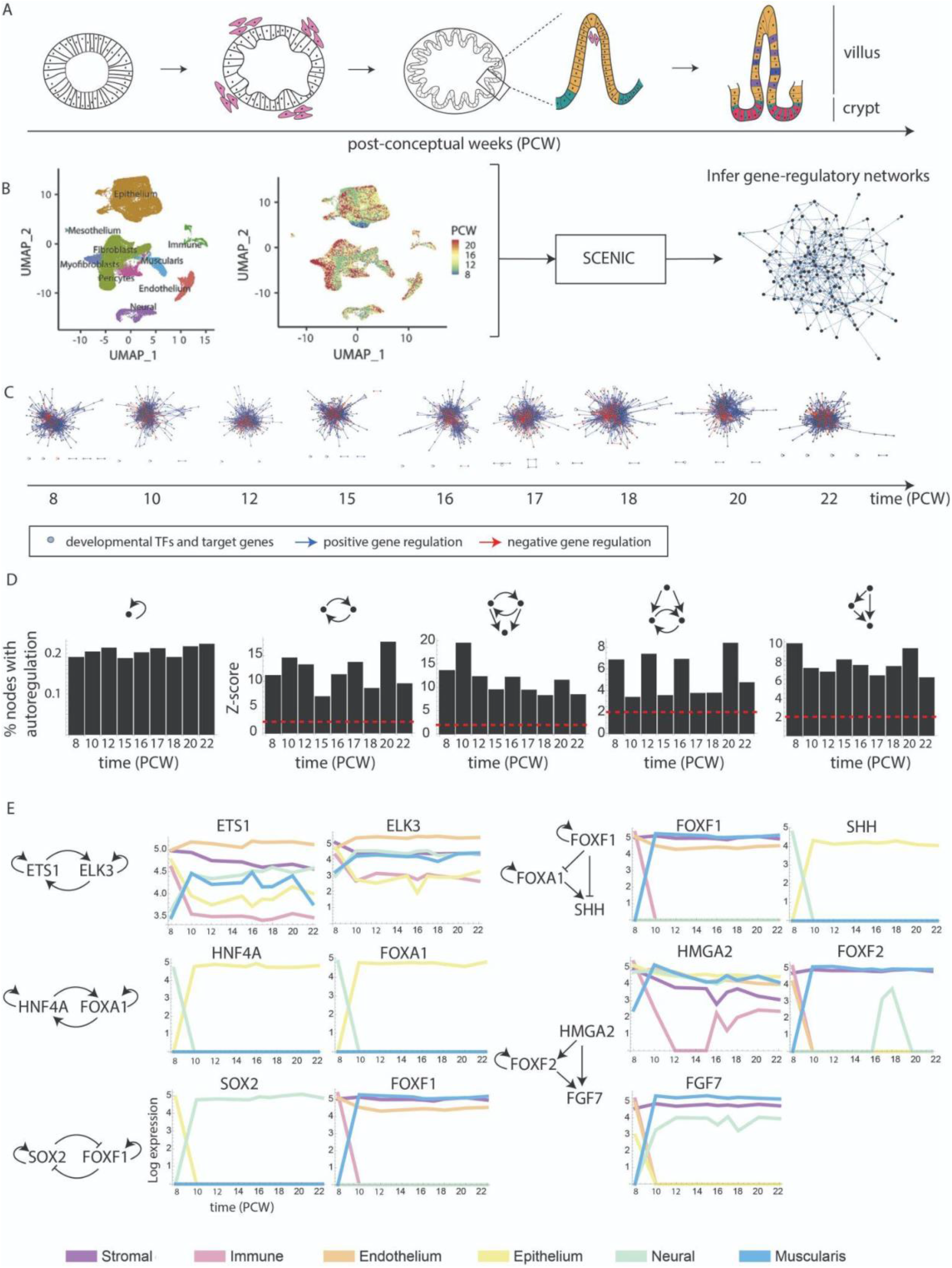
Five network motifs enriched in gene regulatory networks of developing intestine. (A) Schematic of intestine development that is spanned by the data in Fawkner-Corbett et al.^44^. (B) We analyze the single-cell data and use SCENIC to infer gene regulatory interactions between developmental-related TFs and target genes. (C) The GRNs inferred from the data across 9 developmental timepoints. Nodes are TFs and target genes, edges represent positive (blue) and negative (red) interactions. (D) The most statistically enriched network motifs are shown with their Z-score. The red dashed line is a threshold for enrichment significance. For the autoregulation motif (left-most panel) the fraction of nodes in the network that are autoregulated is plotted. (E) Examples of network motifs inferred from 8 PCW. Log mean expression of genes in 6 major cell type categories is plotted as a function of time (PCW).

We find five network motifs (up to 3-node circuits, Methods) that are continuously enriched across all time points. The network motifs are: (1) autoregulation (mostly positive), (2) mutual feedback loop, (3) a regulated feedback loop, (4) a regulating feedback loop, and (5) a feedforward loop (FFL) (Fig 2D). These network motifs are also found in developmental programs of other organisms^40^. The dynamical properties of these 5 network motifs, including bi-stability and delayed response time, are crucial for developmental processes^46,47^. The network motifs we find in intestinal developmental programs can give rise to diverse gene expression patterns from ubiquitous genes expressed in all major cell types to genes expressed in specific cell types (Fig 2E).

### Developmental genes can be categorized based on their position in the network motifs

To explore patterns in the network structure over time, we categorize the genes at each time point according to their position in the network motifs (or their motif role, Fig 1B) and explore whether genes change their positions across time or keep the same motif roles. Considering the 5 network motifs we find in the intestinal developmental programs, there are 7 unique motif roles that we partition the genes into. These motif roles are: autoregulated TFs; TFs in mutual feedback; input, intermediate, and output of an FFL; input to a regulated feedback; and output of a regulating feedback (Fig 3A). Categorizing the genes into these 7 motif roles, we find that the most abundant role in each network is the FFL’s output (Fig 3B). Considering the proportions of genes that generally participate in either input, intermediate or output roles show that intermediate genes form a much smaller group than input and output genes, suggesting a bowtie structure in the networks^48^.

**Figure 3:**
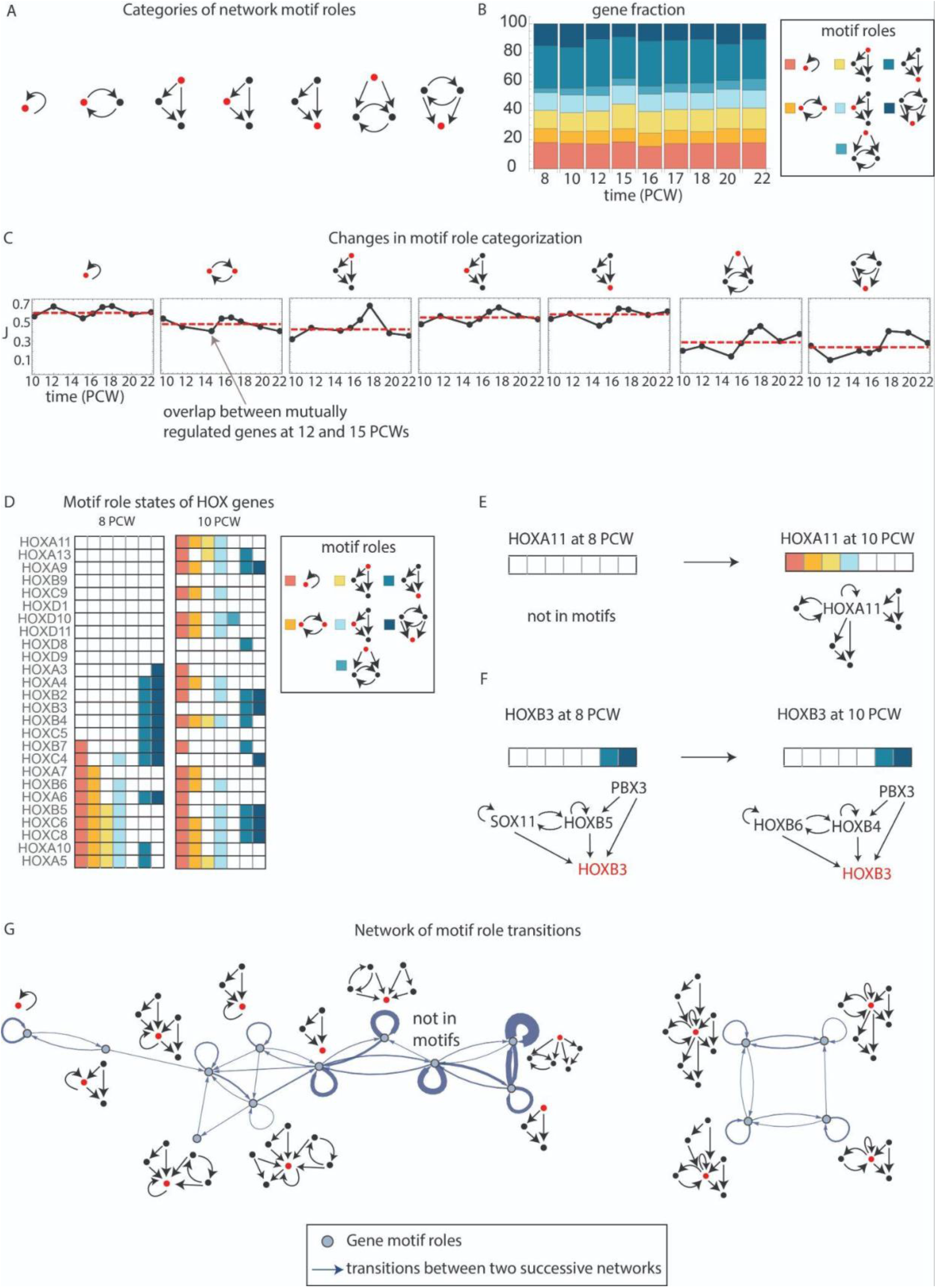
Categorizing developmental genes by motif roles highlights developmental transitions. (A) Developmental genes can be categorized to 7 motif roles (highlighted in red). (B) Fraction of genes participating in each motif role at each time point. (C) In each panel the black points are the Jaccard index between the two sets of genes participating in a particular motif role from two successive time points. The red dashed line is the median Jaccard index across all time points. (D) Table of motif role states of HOX genes at 8 and 10 PCWs. (E) HOXA11 transitions from not participating in motifs to a node common to multiple network motifs combined. (F) HOXB3 maintains the same motif role state between 8 and 10 PCWs despite other genes in the motif that change their regulatory roles. (G) Network of motif role transitions. Nodes in the network represent motif role states and edges represent transitions between two successive networks. Only transitions that occur more than 2 times are considered.

Next, we explore the robustness of categorizing genes according to their role in motifs by looking for changes in the identity of the TFs and genes participating in each motif role. To do so, we compute the overlap (Jaccard index, Methods) between the lists of genes participating in a certain motif role from two successive time points (Fig 3C). We find that autoregulation is the most robust motif role, where about 60% of the TFs that are autoregulated at a certain time point remain so in the next time point in the data. Other motif roles such as the input to a regulated feedback and the output of a regulating feedback are more variable where on average only 30% of the genes participating in these roles maintain their role in successive time points (Fig 3C).

This analysis highlights time points where there are major changes in the identity of genes that participate in a certain motif role. We quantitatively define major changes by considering time points where the Jaccard index is smaller than the median across all time points (red dashed line in Fig 3C). For example, the largest changes in the regulated feedback’s input occur at 10,12, and 15 PCWs (second panel to the right, Fig 3C).

Next, we analyze the TFs and genes that are added in those transition time points to a certain motif role to explore whether these genes are indeed associated with specific expression patterns or are thought to be important for the differentiation of a new cell type. Among the genes added at 10 PCW, we find BMP2, BMP4, and BMP5, which were reported to be expressed in pre-cryptal S2 fibroblasts (telocytes) and to promote epithelial proliferation and differentiation^44^. We also find FOXD1 that controls the terminal ileum, TCF21 which is important for fibroblast early development, and TWIST2 that is involved in key TF networks delineating muscularis cells. FOXF2 which is important for muscle cell differentiation appears in a new motif role at 12 PCW. At 15 PCW, we find several epithelial-restricted genes including LEFTY1 and IHH which are expressed in an epithelial-specific module. At 16 PCW we find WNT5B, which is expressed by stromal, neuronal and muscle cells and supports epithelial cells. At 20 PCW we find FOXL1 which is a marker for S2 fibroblast differentiation, and WNT2B which is expressed in a myofibroblast module. At 22 PCW we find SOX13 which is important for intestinal angiogenesis^44^.

Next, we asked whether changes to gene motif roles across time are random or follow certain rules. To answer this, we define a gene state based on its motif roles (hereby called motif role state) where a gene can participate in one or multiple roles simultaneously (forming a hyper-motif as discussed next). For each gene participating in network motifs at least once, we follow how its motif role state changes over time.

For example, considering HOX genes in our inferred networks (Fig 3D), we find that some HOX genes maintain the same motif role state overtime (Fig 3F) and some transition into new roles (Fig 3E, Fig S2). To visualize the possible transitions between motif role states, we build a network of motif role state transitions (Fig 3G). In this network, the nodes represent a motif role state, and edges represent transitions between states from two successive time points. The weights of the edges are based on the frequency of the transitions. Exploring the patterns of motif role transitions, we find that there are specific rules regarding which role a specific gene can transition into based on its current role within the network. For example, we find that input genes do not transition directly into output genes and vice versa. We also find that TFs that participate in mutual feedback are usually found to be hubs, where they serve as common nodes of multiple network motifs (Fig 3G).

### Network motifs as predictors of gene expression profile

To assess the role of the network motifs we find in developmental programs in regulating gene expression, we next explore the relationship between the identity of network motifs and similarity in gene expression profiles. To do so, we focus on the epithelial and fibroblast cell type populations and consider the specific cell types epithelial and fibroblast progenitor cells diversify into (Fig 4A-B). Within each cell type category, we calculated the Euclidean distance between every pair of cell types (e.g. secretory progenitors and goblet cells, proximal enterocytes and proximal stem cells, etc.) in gene expression space considering their average gene expression profiles across the entire transcriptome (Methods). We then compare this gene expression distance measure to the similarity in the network motifs that we inferred regulating these cells’ gene expression. We estimate the similarity of network motifs by computing the overlap of lists of genes participating in or regulated by a specific network motif between every pair of cells (Methods). We find a large correlation between gene expression distance and overlap in network motif genes where the more network motifs two cell types share, the more similar their gene expression profiles are (Fig 4C-N). This correlation is remarkable given the fact that examining whether a group of ∼20 TFs are autoregulated or not, provides a strong prediction on the similarity in the gene expression state of ∼20,000 genes (Fig 4C, I). We find that some network motifs are better predictors than others. For example, for epithelial cells, autoregulation is more correlated with gene expression than mutual feedback and feedforward loops (Fig 4E,G). However, focusing on effector genes - the identity of genes that are directly regulated by the motifs, improves the correlation (Fig 4D,F,H). Additionally, we find that fibroblast cell types show larger correlations than epithelial cells across all network motifs and their effector genes we examined.

**Figure 4:**
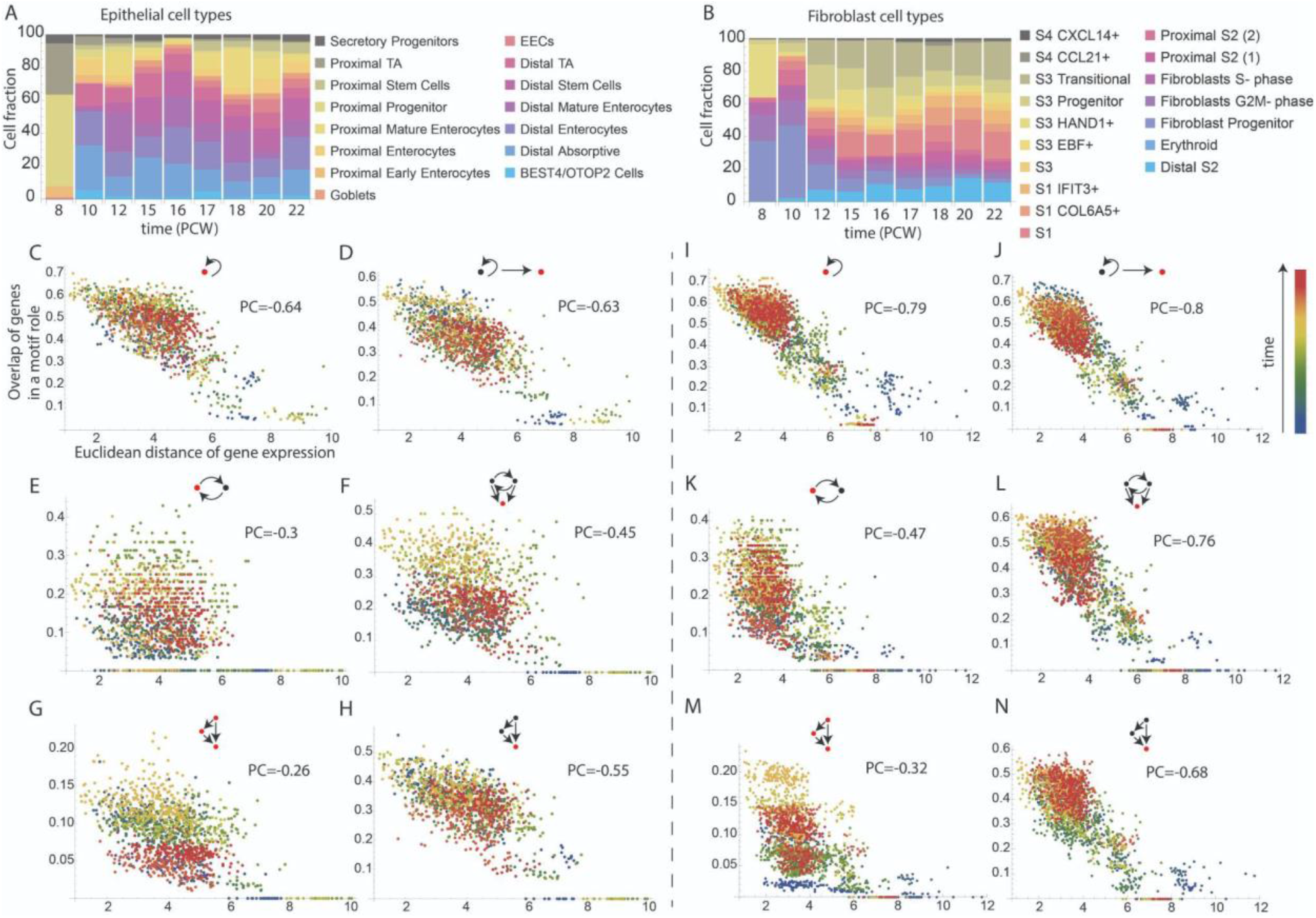
High correlations between network motif identity and gene expression distance in epithelial and fibroblast cell types. (A-B) Cell fraction of epithelial (A) and fibroblast (B) cell types across time. (C-N) Every point is a pair of epithelial (C-H) or fibroblast (I-N) cell types, where the y axis is the overlap (Jaccard index) between genes participating in a network motif or its output gene in the two cell types and the x axis is the Euclidean distance between the gene expression profiles of the two cell types. The overlap in genes is computed for the nodes marked in red in each panel. The color of the points in the plots represents the developmental stage from early stage (8 PCW - blue) to late stage (22 PCW - red). PC=Pearson correlation.

### The recurrent network hyper-motifs show common patterns

We next compute the most statistically enriched hyper-motifs^42^, where multiple motifs are combined through at least one common node (Methods). The most enriched hyper-motifs show several common patterns. Autoregulation is usually coupled to feedback circuits or to FLLs through their intermediate and output nodes. Feedback circuits are combined with FFLs through their intermediate nodes. Motifs share a common input or output node (Fig 5A).

**Figure 5:**
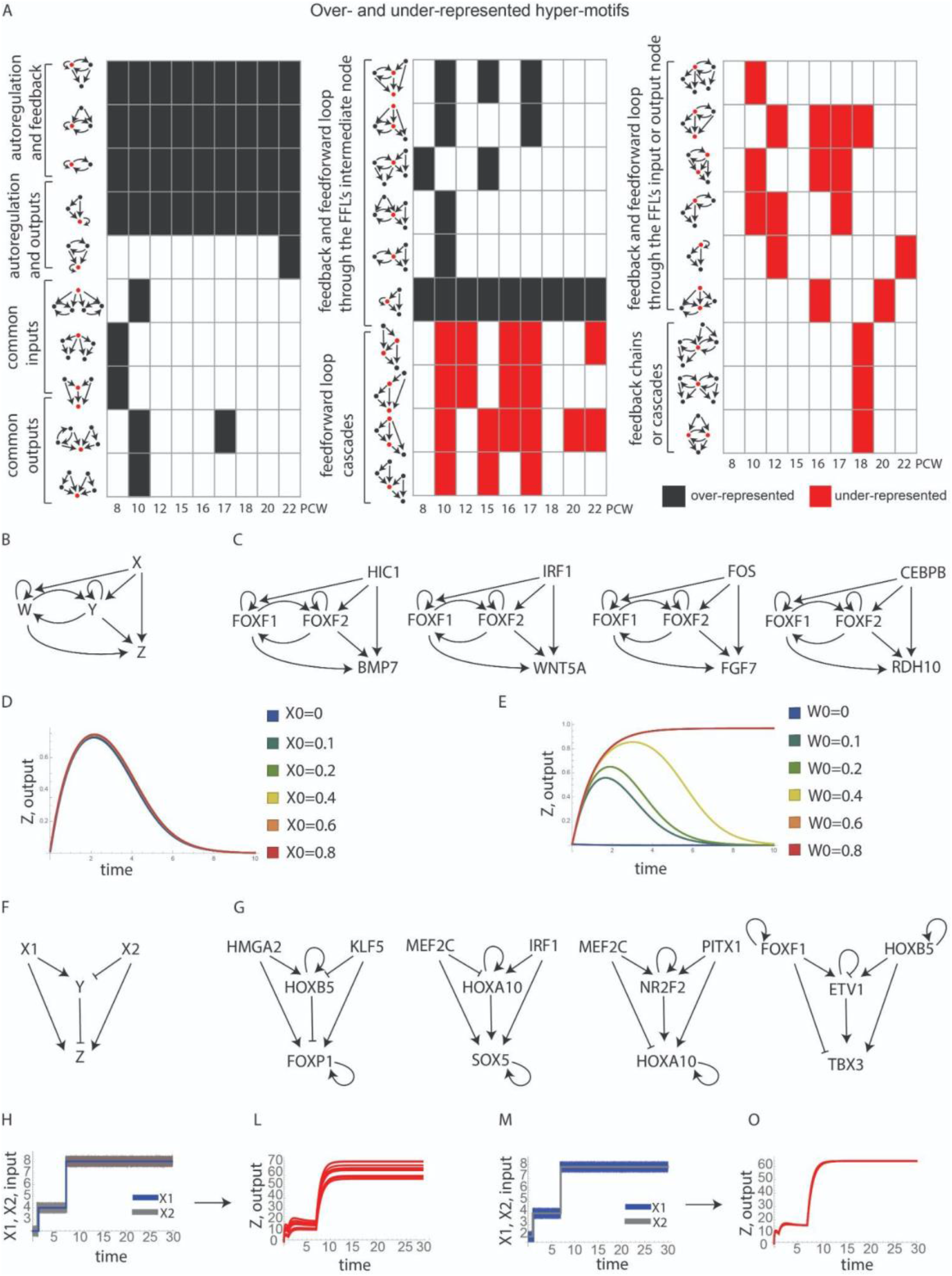
Enriched network hyper-motifs and their dynamical behavior. (A) Table of over- and under-represented hyper-motifs across the developmental process. Rows are different hyper-motif topologies; columns are different time points in the data. Black squares mark positive enrichment, and red squares mark negative enrichment. (B) An enriched hyper-motif circuit where input signal X activates output gene Z and the mutually regulating TFs Y and W which both activate Z as well. (C) Examples of the hyper-motif in the developmental networks. (D-E) Dynamical behavior of the output gene Z where the different curves are for different initial conditions of the input signal X (D) or the TF W (E). (F) Hyper-motif circuit that combines a coherent FFL and an incoherent FFL. (G) Examples of the hyper-motif in the developmental networks. (H) Simultaneous step-like changes in X1 (noisy, gray) and X2 (precise, blue) (L) Dynamical behavior of output Z in response to the input signals plotted in H. (M-O) Same as H-L only with X1 as the noisy signal and X2 as the precise signal.

Some combinations are statistically enriched across all developmental time points, while others are significantly enriched only at certain time points. Assuming that some combinations of motifs may lead to new properties, time points with enrichment of hyper-motifs suggests emergence of new cell types or tissue-level patterns. The most enriched hyper-motif topologies appear at 10 PCW, which is indeed the time where the crypt to villus axis is formed (Fig 5A).

The under-represented network hyper-motifs also follow certain patterns. Autoregulation and mutual feedback do not combine with the FFL’s input node, combinations of multiple feedback circuits are avoided only at 18 PCW, and cascades of FFLs are also excluded in multiple time points (Fig 6B).

**Figure 6:**
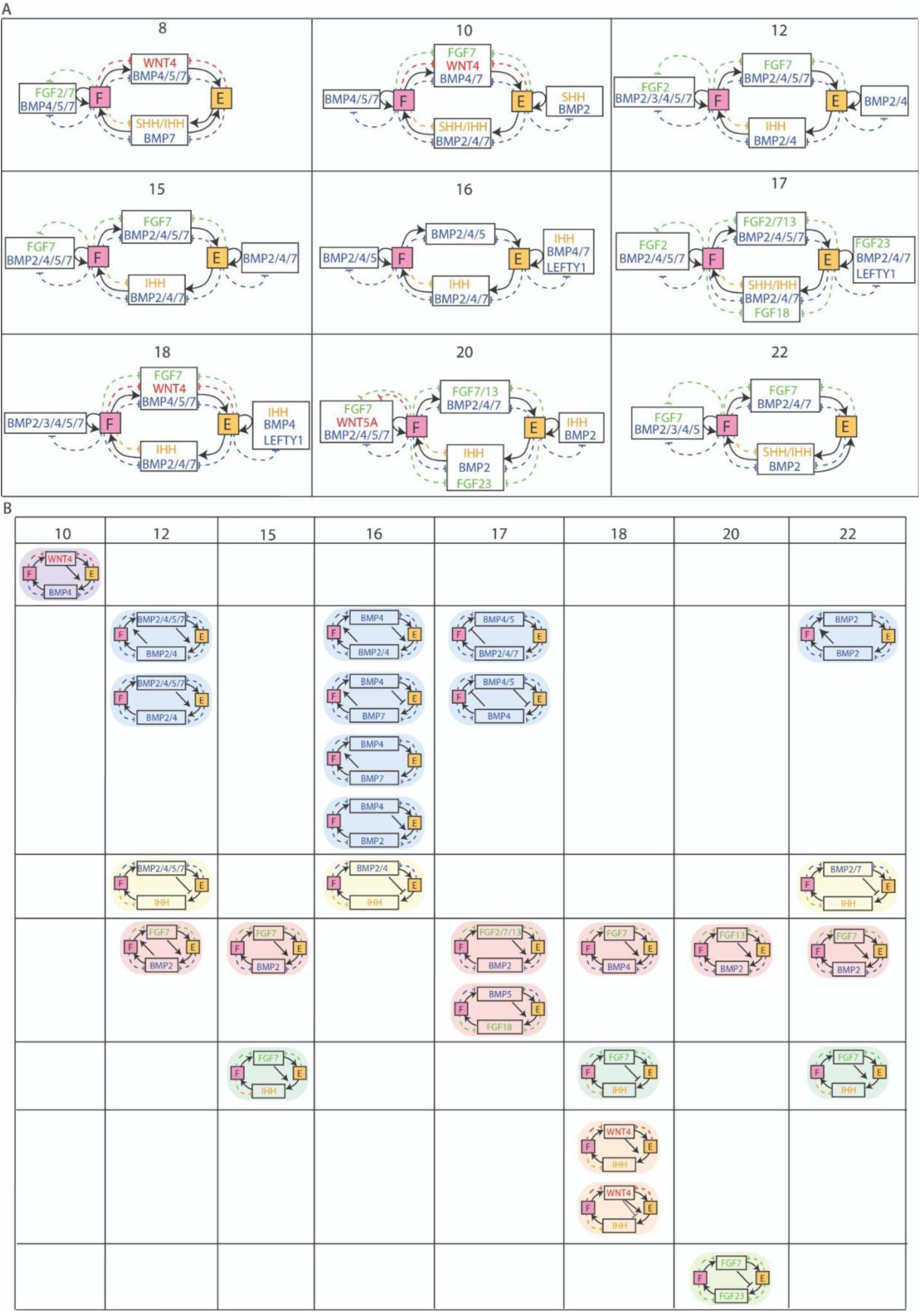
Inferring hyper-motif circuits between communicating cells through ligand-receptor interactions. (A) Cell-circuits between the epithelial (E, orange square) and fibroblast (F, pink square) cell type populations through morphogen and growth factor (GF) provision from 8-22 PCWs. Black arrows from cells to GFs represent ligand production. Black arrows from GFs to cells represent binding to cognate receptors. Dashed colored inhibitory arrows represent inhibition of GFs through antagonist expression. (B) Table of cell circuits where expression of morphogen ligands is directly regulated by a regulatory circuit downstream to a morphogen signaling pathway. Black arrows pointing from GF1 to another GF’s production arrow, GF2, represent activation or inhibition through a gene regulatory circuit that is downstream to the GF1 signaling pathway in the cell type producing the GF2 signal.

### Dynamical properties of enriched hyper-motif regulatory circuits

We consider several hyper-motif regulatory circuits that we find to be statistically enriched in the networks and mathematically model their dynamical behavior. One of the enriched hyper-motifs is a combination of a feedback circuit and an FFL through the intermediate node of the FFL (Fig 5B). This hyper-motif is very frequent in the regulatory networks of stromal and epithelial cells throughout the developmental process where it directly controls the expression levels of morphogen ligands and receptors as well as various development-related TFs (Fig 5C). We developed a mathematical model for the dynamical behavior of this hyper-motif (Methods). Our model suggests an interesting property of this hyper-motif where there are two types of input signals affecting the output gene. The first input signal is the FFL’s input X, and the second class of input signal are the Y/W TFs which mutually regulate each other. While both inputs are assumed to be required for the transcription of the output gene (Z) in our model, they differ in their effect on the dynamical behavior of Z. The X input signal is required to turn on Z, but its initial or final levels do not influence the dynamical behavior of Z. This means that Z is insensitive to variations in the levels of X (Fig 5D). In contrast, the initial levels of Y or W have a large effect on the dynamical behavior of Z (Fig 5E). This sensitivity to the Y/W levels provides a way of generating different responses to varying levels of inputs using a simple regulatory circuit, thus fulfilling the requirement for the interpretation of morphogen gradients, for example.

We also consider a hyper-motif which joins two types of FFL circuits: coherent and incoherent FFLs (Fig 5F). This circuit is seen for example at 8 PCWs in fibroblast cells where the HMGA2-HOXB5-FOXP1 circuit is an incoherent FFL (HMGA2 activates FOXP1 directly but inhibit it through HOXB5), whereas the KLF5-HOXB5-FOXP1 circuit is a coherent FFL (Fig 5G). This hyper-motif can exhibit known properties of each motif when the input signals vary separately. When the two input signals are changing simultaneously, this motif presents an interesting behavior where the final level of the output Z is robust to variations in the input signal regulating the incoherent FFL (X1) but is sensitive to noisy inputs stimulating the coherent FFL (X2) (Fig 5H-O).

### Cell-cell hyper-motif circuits emerge through morphogen signaling pathways

Next, we explore how regulatory circuits within cells are linked through morphogen signaling pathways creating hyper-motif circuits between communicating cells. For the purpose of cell communication circuit analysis, we focus on fibroblast and epithelial cell type populations, the two cell types playing important roles in governing intestinal development^49^. First, we identify pairs of interacting cells where we consider interactions through ligand-receptor pairs of the developmental morphogen families (BMP, WNT, HH, FGF, and retinoic acid). To identify pairs of interacting cells, we used NicheNet, a computational method that infers ligand-receptor interactions considering expression patterns of ligands, receptors and downstream signaling genes in the receiving cells^50^ (Methods). Using NicheNet, we highlight the top ligands that are predicted to link fibroblast and epithelial cell types (Fig 6A). Throughout the developmental process, we find that fibroblasts and epithelial cells communicate reciprocally through various BMP ligands; whereas IHH and SHH production is restricted to epithelial cells and its receptors and antagonists are produced by fibroblasts. Other morphogens, such as WNTs, appear at specific time points (Fig 6A).

Next, we use these cell communication circuits to infer regulatory networks linking perception of certain morphogens to production of other morphogens. We focus on reciprocal circuits composed of provisions of pairs of signaling pathways. Out of these circuits, we look for situations where gene regulatory interactions that are downstream to a certain morphogen signaling pathway in the receiving cell directly upregulate or downregulate expression of a morphogen ligand (Fig 6B). Following these morphogen circuits over time, we find several patterns that recur in multiple time points. For example, downregulation of IHH by BMP signaling appears at 12, 16, and 22 PCWs.

### Cell-cell hyper-motif circuits can provide emergent dynamical properties

We consider three examples of hyper-motif regulatory circuits between communicating cells and examine their potential dynamical behavior in more detail. To do so, we consider the shortest path in the gene regulatory network linking the TF (Y) downstream to the input morphogen signaling pathway (X) received by the cell and the output morphogen ligand (Z) that is produced by the same cell (Fig 7A). Since the genes participating in this shortest path are part of a larger complex gene regulatory network, we developed a new method to extract the circuit topology controlling the expression of the output gene Z. Instead of simply considering a cascade of genes following the shortest path between X and Z in the network, we are inferring the ‘*shortest network motif path*’. Using the shortest network motif path approach, we follow the network motifs in which the genes linking X and Z participate in (Fig 7A, Methods). After inferring the shortest network motif path between a pair of morphogen signals, we developed a mathematical model to simulate the dynamical behavior of the resulting hyper-motif circuit (Methods). We find that interactions between members of BMPs and IHH (Fig 7B) can provide pulsatile response (Fig 7C) and pure oscillations (Fig 7D). Interactions between BMP4, WNT4 and its inhibitor SFRP2 (Fig 7E) can yield antagonistic expression patterns for BMP4 and WNT4 depending on model parameters (Fig 7F-G). The hyper-motif in epithelial cells that regulates IHH and is activated by WNT4 (Fig 7H) can provide pulsatile (Fig 7I) or delayed and sustained (Fig 7J) response.

**Figure 7:**
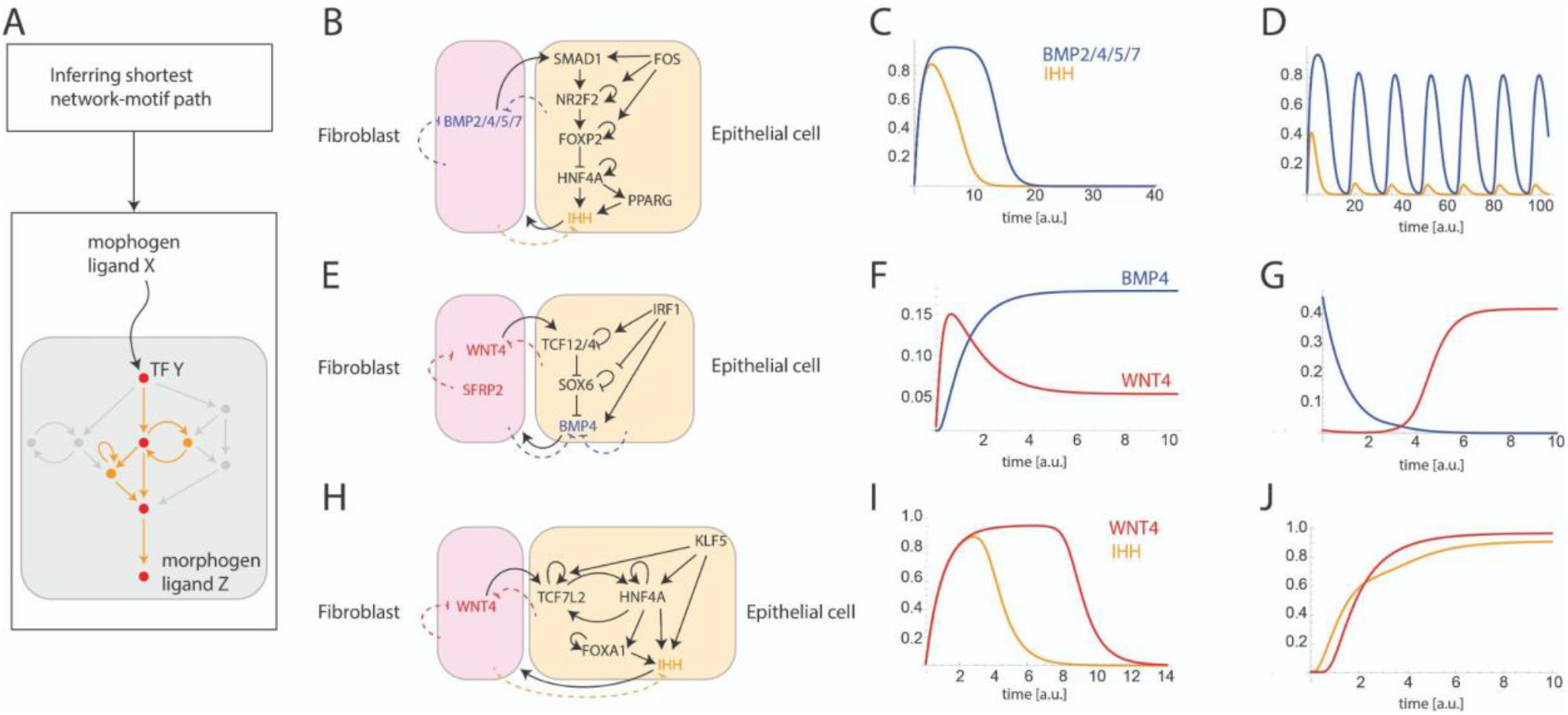
Dynamical properties of cell-cell hyper-motif circuits. (A) Illustration of our method to infer the shortest network motif path linking input signal X and output gene Z. Nodes participating in the shortest path are marked in red. Edges and additional nodes participating in the shortest network motif path are marked in orange. (B-J) Examples of three hyper-motif gene regulatory circuit topologies (B, E, H) linking two morphogen signaling pathways between fibroblast and epithelial cells. Dashed arrows represent production of morphogen antagonists. (C-D, F-G, I-J) Simulated dynamical behavior of the circuits in B, E and H with varying parameters or initial conditions (Methods).

### Hyper-motif regulatory circuits frequently regulate morphogen production

Next, we explore the extent to which hyper-motif regulatory circuits regulate morphogen expression directly through regulation of morphogen ligands and indirectly, through regulation of the morphogen receptors and antagonists. For every pair of morphogen families, we examine whether there is a direct path in the gene regulatory network linking one morphogen signaling pathway to expression of another morphogen’s ligand, receptor or antagonist. Counting the number of target genes belonging to a certain morphogen family that are regulated by another morphogen signaling show large variation between the pairs of morphogens, as well as variation in the pattern over time (Fig 8A, C). Additionally, we explore the number of genes, within the regulatory paths linking pairs of morphogen signaling, that participate in network hyper-motif circuits. We find that most regulatory paths linking morphogen pairs are enriched with hyper-motif regulatory circuits (Fig 8B, D). Examining the composition of the most common hyper-motif regulatory circuits within these paths, we find that many of the regulatory circuits regulating expression of morphogen signaling correspond to the most statistically enriched hyper-motifs in the network (Fig 8E-F; Fig 5A), suggesting that these enriched hyper-motif circuits indeed play important roles in regulating morphogen expression patterns.

**Figure 8:**
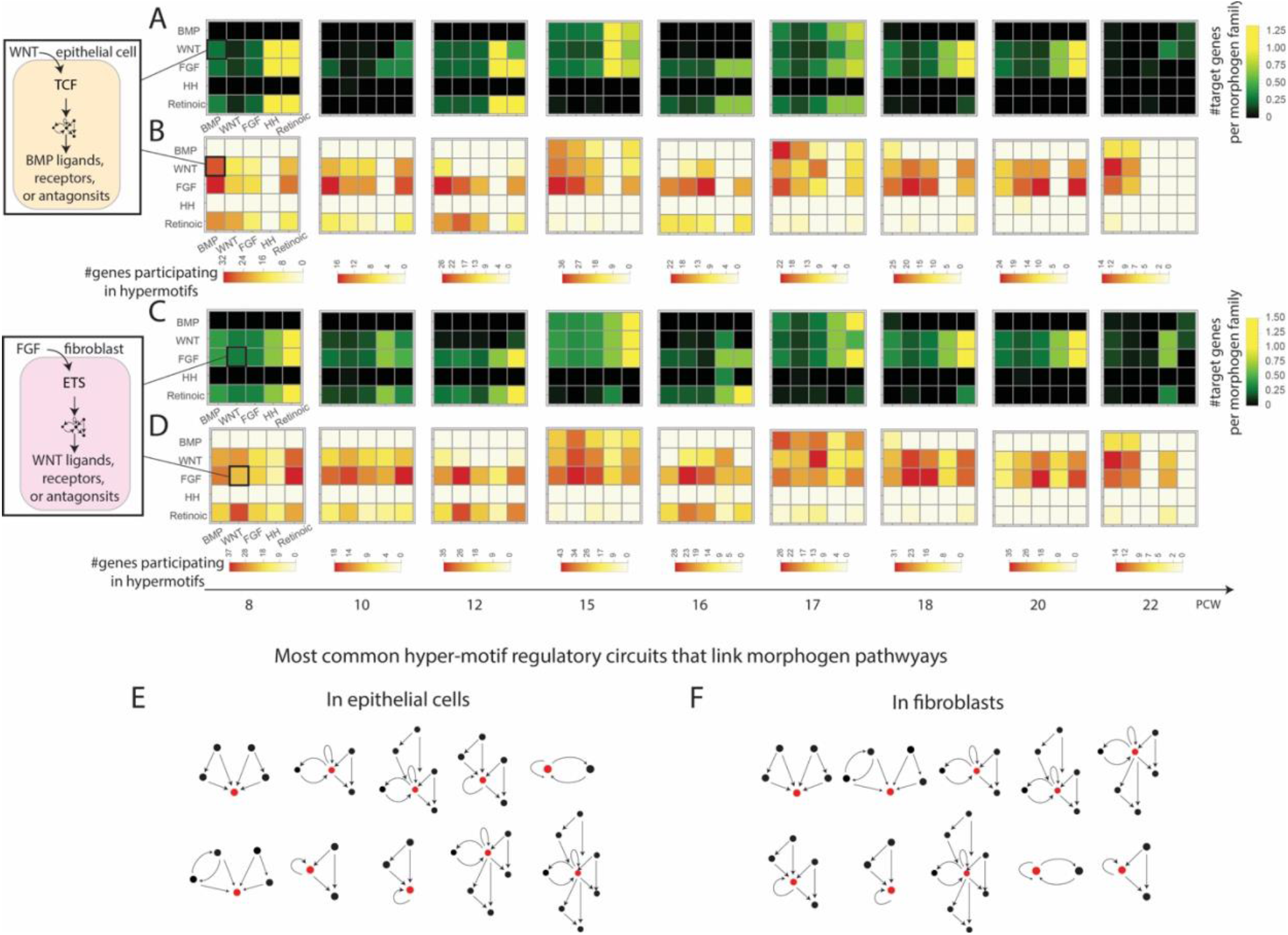
Role of hyper-motif circuits in regulating morphogen signaling pathways. (A, C) Heatmaps of the number of target genes that are regulated downstream to morphogen signaling pathways in epithelial cells (A) and in fibroblasts (C). A square in X row and Y column is colored by the number of ligands, receptors and antagonists belonging to morphogen family Y that are found to be directly regulated in response to morphogen ligands from the X family, divided by the total number of ligands in the Y family. (B, D) Heatmaps of the number of genes in the regulatory circuits described in (A) and (C) that participate in more than one motif role (forming a hyper - motif) in epithelial cells (B) and fibroblasts (D). (E-F) The 10 most common hyper-motifs regulatory circuits linking morphogen pathways in epithelial cells (E) and fibroblasts (F).

## Discussion

Developmental processes are regulated by complex regulatory networks. To disentangle this complexity, we used our recently developed framework exploring how building-block circuits are joined in complex networks to reveal design principles of developmental programs. By applying our approach to intestinal development single-cell data, we found that developmental GRNs show 5 network motifs that are joined in the network according to specific rules. We find that network motif identity and position of genes within motifs are good predictors for gene expression profiles for certain network motifs within epithelial and fibroblast populations, supporting the validity of the network motif approach to examine gene regulatory networks. Examining major transitions in the network structure, including emergence of enriched network hyper-motifs and appearance of new genes in key positions within the network, highlight major developmental events and their potential regulators. Lastly, we find that morphogen ligands, receptors and co-receptors are regulated directly by combinations of network motifs creating hype-motif circuits between communicating cells that can generate emergent dynamical properties.

Our approach offers a new way to categorize developmental genes according to their roles in the network motifs. This categorization distinguishes between input genes, effector genes and hub genes that are core regulators combining multiple network motifs. It will be interesting to apply this method to developmental processes of other tissues to explore the robustness of gene categorization based on network motif roles.

In our analysis of over- and under-represented network hyper-motifs, we discovered that positive feedback circuits, including autoregulatory and mutual feedback, are highly enriched throughout the developmental process, whereas cascades of feedforward loops are statistically scarce. This finding is surprising, given that both circuit designs provide a delayed response which is an important property in developmental programs. The potential selection against cascades of FFLs may be due to the fact that FFL cascades provide delay in response time at the cost of noise amplification, whereas positive feedback circuits provide delay and sensitivity to input signals without compromising on the ability to buffer noise propagation^51,52^.

We note that our approach provides theoretical predictions on the mesoscale structure of developmental gene regulatory networks at a systems level. Future work where predictions regarding particular TF and target gene positions and roles within the networks are experimentally explored will be integral to complement this approach.

## Acknowledgments

We would like to thank Eviatar Weizman from the Mantoux Bioinformatics institute of the Nancy and Stephen Grand Israel National Center for Personalized Medicine, Weizmann Institute of Science, for providing support in the application of the NicheNet approach. We would like to thank Scott Pope for supporting initial TF binding motif analysis. We thank all current and former members of the Medzhitov lab as well as David Fawkner-Corbett and Agne Antanaviciute for discussion on this project.

## Methods

### Inference of gene regulatory networks from single-cell data

We analyzed single-cell RNA sequencing data from Fawkner-Corbett et al.^44^ using the SCENIC package in python (pySCENIC)^45,53^. The pySCENIC pipeline consists of: (1) Obtaining expression data per each time point in the data using the Seurat pipeline^54^; (2) Inferring an adjacency matrix describing co-expression relationships between transcription factors (TFs) and target genes; and (3) Constructing regulons - modules containing TF and target genes by integrating motif information. The activity of a TF in a cell is calculated by integrating the TF target genes’ expression. We repeated the pySCENIC pipeline 6 times and kept the overlapping regulons to reduce noise. To focus on TFs with variable activity in the tissue, we filtered out low-variance regulons by removing those with activity variance less than 0.1. To build the gene regulatory networks, we considered pairs of TFs and target genes that have a predicted activity that is higher than the median across all pairs, considering genes that belong to a manually curated list of developmental-related genes (5143 genes).

### Network motif enrichment analysis

To identify network motifs - statistically enriched patterns in the developmental regulatory networks, we used the MFinder package^55^ to identify network motifs of up to 3 nodes. We verified these network motifs by detecting enriched patterns using the IGraphM package in Mathematica. To randomize the networks, we used the IGRewire function and to find motifs and their frequencies we used the IGMotifs function.

### Motif role transitions

To explore transitions of genes between network motif roles, we first categorized the genes in the regulatory networks according to their role (or roles) in the motifs. We denote *N*_*m,t*_ as the set of nodes in a GRN that participate in network motifs in time t. Next, for each time point t, we categorized nodes in *N*_*m,t*_ to 7 groups according to their roles in the network motifs: {*n*_*1,t*_, *n*_*2,t*_, · · ·, *n*_*7,t*_}, for the 7 unique motif roles in our case (Fig 3A). Note that certain nodes may belong to more than one group (which is the case where network motifs are combined). Next, for each one of the 7 motif roles, we consider all successive pairs of node groups within each role, such as: {*n*_*1,t*_, *n*_*1,t*+*1*_}, {*n*_*1,t*+*1*_, *n*_*1,t*+*2*_}, etc. For each pair of successive groups, we compute the Jaccard index: *J*(*n*_*i,t*_, *n*_*i,t*+*1*_) = |*n*_*i,t*_ ∩ *n*_*i,t*+*1*_|/|*n*_*i,t*_ ∪ *n*_*i,t*+*1*_|, which is the size of the intersection of the two groups divided by the size of their union. If all genes in a certain motif role remain in the same motif role in the next time point: *J* = *1*, and if none of the genes remain in the same motif role: *J* = *0*.

To build the network of motif role transitions, we defined motif role states for every gene (g) in the networks in each time point (t): *x*_*g,t*_, which is a 7-dimensional vector that represents whether gene g participates in each one of the 7 motif roles or not (for example: (*1,0,0,0,0,0,0*), (*1,1,0,0,0,0,0*), (*1,0,1,0,0,0,0*), etc.). Overall, there are *2*^*7*^ = *128* different possible motif role states (although not all of them are implemented in the developmental GRNs). We then build a network where the nodes represent the different possible motif role states. We then consider for every gene, all pairs of motif role states from successive time points: {*x*_*g,t*_−> *x*_*g,t*+*1*_}. For each direct transition between motif role states, we draw an edge connecting the two motif role states (or a self-loop arrow in case the genes retain the same motif role state). We assign weights to the edges to account for the observed frequencies of each transition.

### Correlations between network motif identities and gene expression profiles

We consider pairs of specific cell types (clusters) within the epithelial cells (and separately also within the fibroblast populations) across successive time points in the data. For each such pair of cell types and for every pair of successive time points, we compute the Jaccard index of genes participating in a certain network motif or are regulated by a certain network motif. For example, considering the feedforward loop (FFL), first we create a list of all the triplets of genes participating in FFLs in each cell type. We then compute the Jaccard index of the two lists of triplet genes. We compare the Jaccard index with the Euclidean distance between the gene expression profiles of the two cell types in question. We then used the PearsonCorrelationTest function in Mathematica to estimate the Pearson correlation for every network motif and its statistical significance.

### Detection of over- and under-represented network hyper-motifs

To detect over- and under-represented network hyper-motifs, we detect enrichment of combinations of network motifs. To do so, we follow the following steps (presented in detail in ^42^):

1. Compute the Jaccard index for all *k*(*k* − *1*)/*2* pairs of motif roles (here *k* = *7*) within each time point: *J*(*n*_*i,t*_, *n*_*j,t*_) = |*n*_*i,t*_ ∩ *n*_*j,t*_|/|*n*_*i,t*_ ∪ *n*_*j,t*_|·
2. For nodes that appear more than once in the same role within the same network, we compute the Jaccard index by the ratio of the number of nodes that appear in a network motif role more than once to the total number of nodes that participate in that motif role.
3. We use the MFinder package^55^ to create 100 random networks (for each time point in the data) that have the same number of nodes and edges, the same number of incoming and outgoing edges per node, and the same frequency of all subgraphs up to three-node subgraphs, as in the real network.
4. After categorizing the nodes according to their motif roles for each of the 100 random networks (per time point), we repeat steps 1 through 2 for all the random networks and compute *J*_*rand*_ (*n*_*i,t*_, *n*_*j,t*_) for all pairs of motif roles in the random networks.
5. For every *i,j* such that *i, j* ∈ {*1,2*, · · ·, *k*}, we compute the Z-score of *J*(*n*_*i,t*_, *n*_*j,t*_) from the distribution of {*J*_*rand*_(*n*_*i,t*_, *n*_*j,t*_)}: *Z*_*ij*_ = (*J*(*n*_*i,t*_, *n*_*j,t*_) − *mean* ({*J*_*rand*_(*n*_*i,t*_, *n*_*j,t*_)}))/ *std*({*J*_*rand*_ (*n*_*i,t*_, *n*_*j,t*_)}), and compute the *P* value by estimating the cumulative density of *Z*_*ij*_ for a normal distribution with zero mean and unit variance. We use the function NormalPValue in the package HypothesisTesting in Mathematica to compute the *P* value. We then use the Benjamini–Hochberg procedure to correct for multiple hypothesis testing, which provides us a corrected q-value for each pair of motif roles.
6. We consider over- and under-represented motif combinations if their q-value is smaller than 0.05, where *Z*_*ij*_ > *0* for over-represented combinations and *Z*_*ij*_ < *0* for under-represented combinations.

### Mathematical models for statistically enriched network hyper-motif regulatory circuits

We consider the following differential equations to model the dynamical behavior of the hyper-motif circuit that combines two FFLs and a mutual feedback loop (Fig 5B):

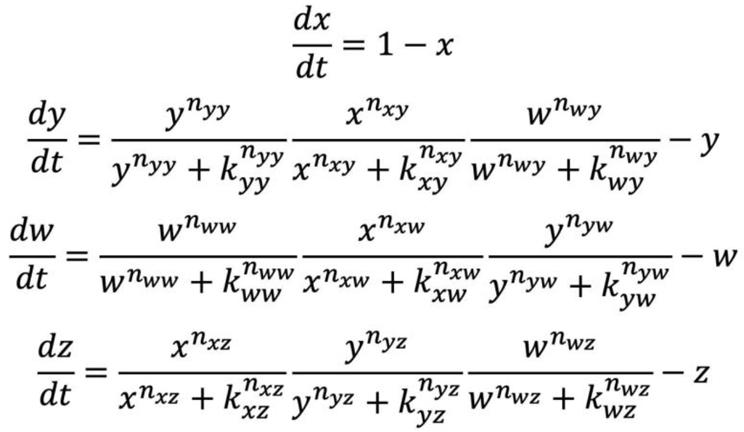

We chose parameter values that provide bistability to the mutual feedback circuit. We therefore used the following parameter values:

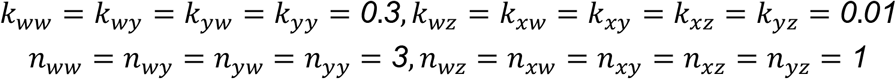

We used the following initial conditions for Fig 5D:

*y*_*0*_ = *0*·*25, w*_*0*_ = *0*·*35, z*_*0*_ = *0*·*01*, and *x*_*0*_ as denoted in Fig 5D. and the following initial conditions for Fig 5E:

*x*_*0*_ = *0, w*_*0*_ = *0*·*3, z*_*0*_ = *0*·*01*, and *y*_*0*_ as denoted in Fig 5E.

We consider the following differential equations to model the dynamical behavior of the hyper-motif circuit that combines two FFLs forming a multi-input FFL (Fig 5F):

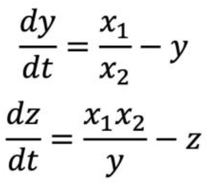

Here we used log linear terms (assuming that the variables are far from saturation) since this is the range of parameters that provide well-known dynamical properties of the coherent and incoherent FFLs such as the fold-change detection property^39,56–58^.

### Inference of cell communication circuits through morphogen ligand-receptor interactions

To infer interactions between epithelial and fibroblast populations, we applied NicheNet, an algorithm that infers ligand-receptor interactions within a transcriptomic dataset, using prior knowledge on ligand-receptor interactions and regulatory networks^50^. We applied NicheNet to the intestinal development scRNAseq dataset considering interactions between the specific cell type populations of fibroblasts and epithelial cells across the developmental process. The pairwise interaction between cell types (mature enterocytes, fibroblast progenitors, absorptive cells, etc.) was calculated independently for each time point in the data (9 time points spanning 8-22 PCWs). For each cell type and for each time point, genes of interest (GOIs) were defined as genes that were upregulated in a particular cell type compared to all other cell types in that time point and expressed in at least 25% of the cells within the cell type population. Interaction score was calculated as the Pearson coefficient calculated by ‘predict_ligand_activities’ function, using the GOIs of that time point and the genes that were expressed in the ‘receiver’ clusters as targets. For each pair of sender-receiver cells, we chose the top 10% quantile ligands. This analysis allowed us to infer interactions between every pair of specific cell type populations among the different fibroblast and epithelial clusters. In Fig 6A we consider the possible interactions through the different morphogen ligand-receptor pairs where we group together the various fibroblast and epithelial clusters.

### Analysis of shortest network-motif paths

To infer regulatory interactions that link a signal (S) perceived by a cell to an expression of a ligand encoded by a target gene (Z), we consider the GRN of that cell type. We then use the function ‘FindShortestPath’ in Mathematica to compute the shortest path linking a transcription factor X, which is downstream to the S signaling pathway and the target gene Z. Let us denote the set of genes that are in this shortest path {*G*_*s*_}. We now build a new set of genes that will make up the shortest network-motif path: {*G*_*M*_}. For each gene *g*_*i*_ in {*G*_*s*_}, we perform the following steps:

1. Check if *g*_*i*_ participates in network motifs in the same time point and in the same cell type.
2. If the answer to #1 is yes - add *g*_*i*_ and the genes interacting with *g*_*i*_ in the network motif to {*G*_*M*_}. *Note that in case *g*_*i*_ participates in multiple network motifs of the same topology and in the same motif role, we consider only one instance of this motif in {*G*_*M*_}. This is similar to having multiple shortest paths in a network, and arbitrarily choosing one of them.
3. If the answer to #1 is no - add only *g*_*i*_ to {*G*_*M*_}.

### Mathematical models for regulatory circuits between communicating cells

We model the dynamical behavior of two morphogen lignands that regulate each other’s expression through regulatory circuits between a pair of communicating cells (Fig 7). For the cell circuit depicted in Fig 7B we use the following set of differential equations where x represents the FOS/SMAD1/NR2F2/FOXP2 module, y represents HNF4A, Z is PPARG, u is for IHH, and v represents BMP2/4/5/7:

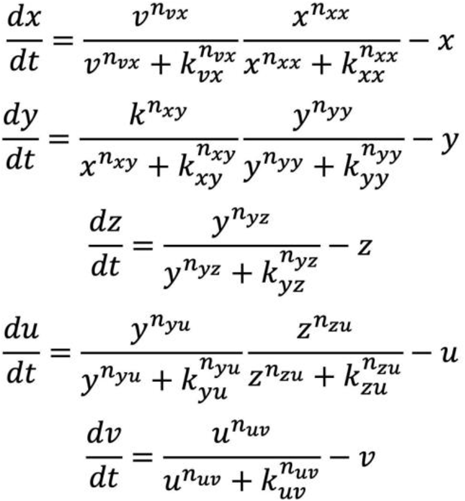

We used the following parameter values for Fig 7C:

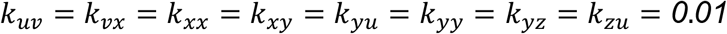

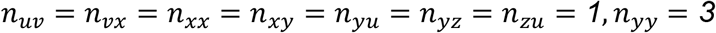

with the following initial conditions:

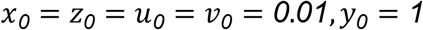

And the following parameter values for Fig 7D:

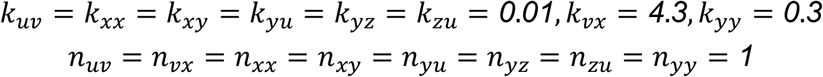

with the following initial conditions:

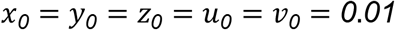

For the cell circuit depicted in Fig 7E we use the following set of differential equations where x is IRF1, y is TCF12/4, Z is SOX6, u is for BMP4, and v represents WNT4. To account for effect of expression of BMP and WNT antagonists, we added direct inhibition terms in the equations for the morphogen ligands:

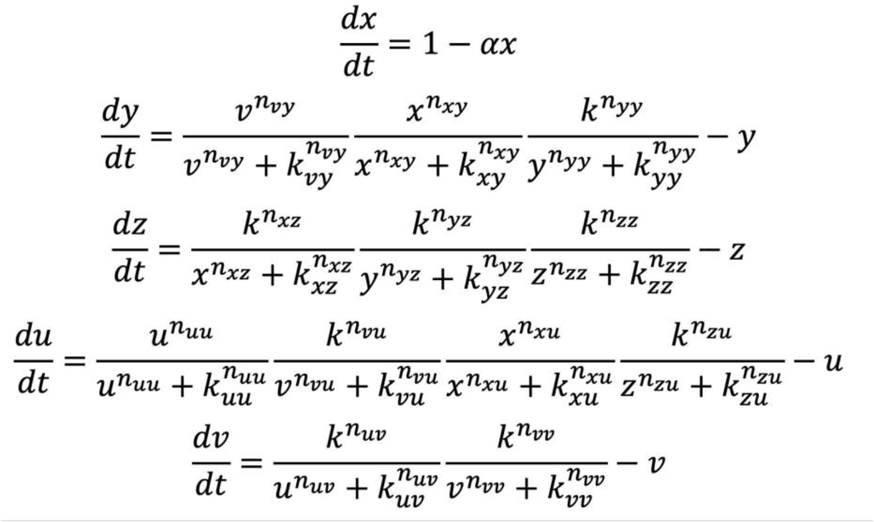

We used the following parameter values for Fig 7F:

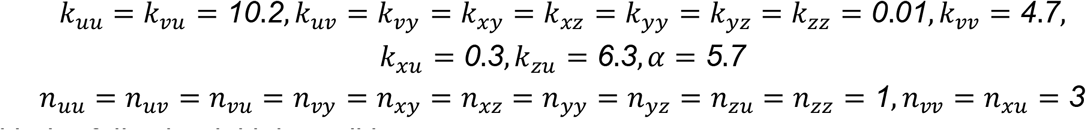

with the following initial conditions:

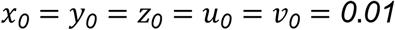

And the following parameter values for Fig 7G:

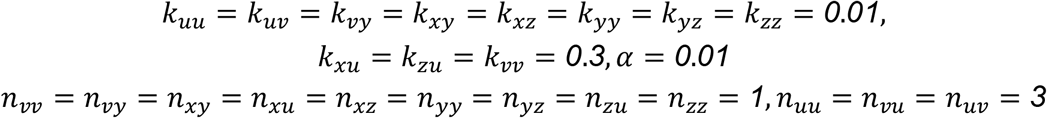

with the following initial conditions:

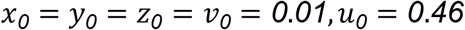

For the cell circuit depicted in Fig 7H we use the following set of differential equations where x is KLF5, y is TCF7L2, w is HNF4A, Z is FOXA1, u is for IHH, and v represents WNT4:

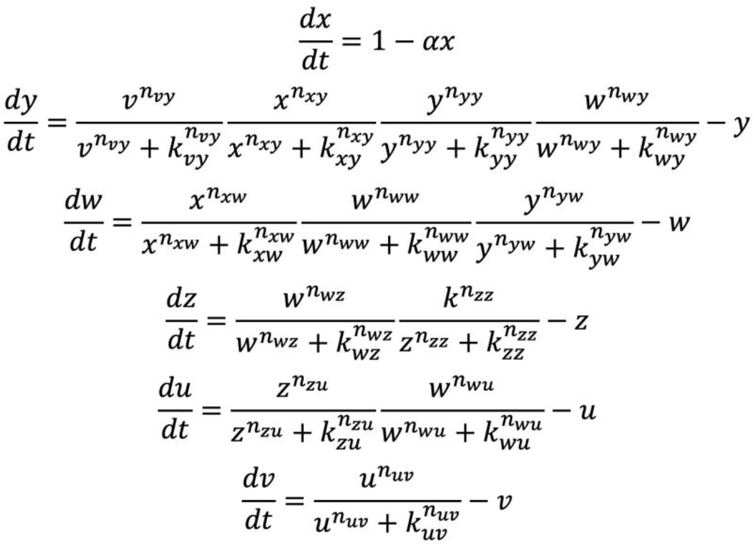

We used the following parameter values for Fig 7I:

*k*_*ij*_ = *0*·*3, n*_*ij*_ = *3* for every i,j with the following initial conditions:

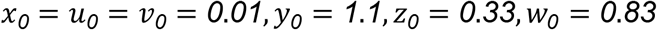

And the following parameter values for Fig 7J:

*n*_*ij*_ = *3* for every i,j except for *n*_*vy*_ = *1*

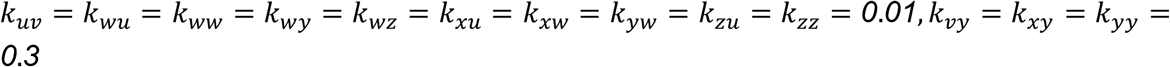

with the following initial conditions:

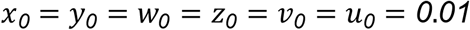

## SI Figures

**Figure S1:**
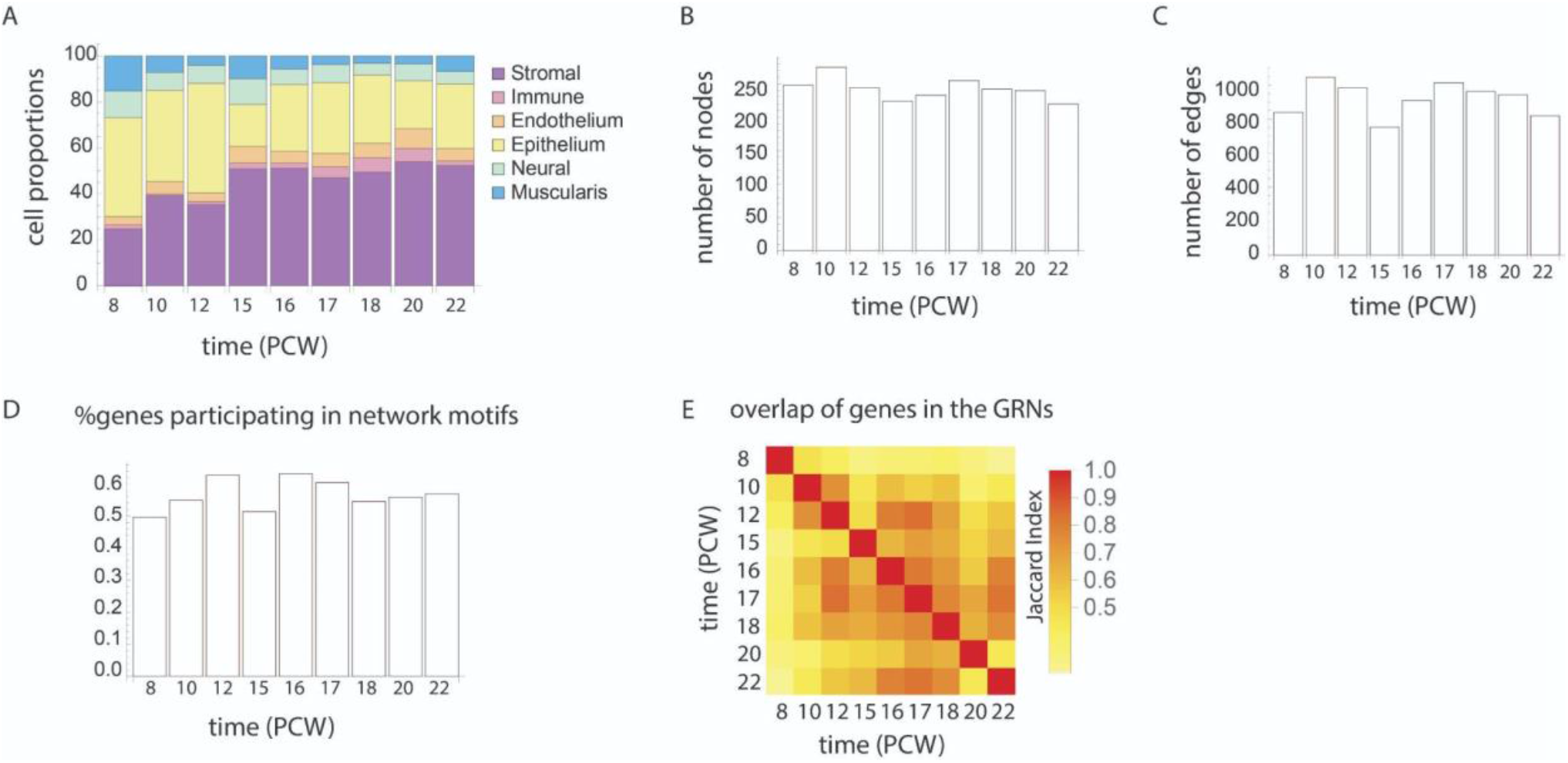
(A) Proportions of the 6 major cell type categories in the intestine computed from the single-cell data in Fawkner-Corbett et al.^44^ across the 9 time points that follow the developmental process. (B-C) Number of nodes (B) and edges (C) in each network inferred from the single-cell data. (D) Proportion of genes that participate in network motifs out of the total number of genes in each network. (E) Heatmap of the overlap of genes that participate in the different gene regulatory networks (GRNs). Every square is a comparison between two networks from time points in the data. Overlap is computed by calculating the Jaccard index of the two lists of genes participating in the two networks.

**Figure S2:**
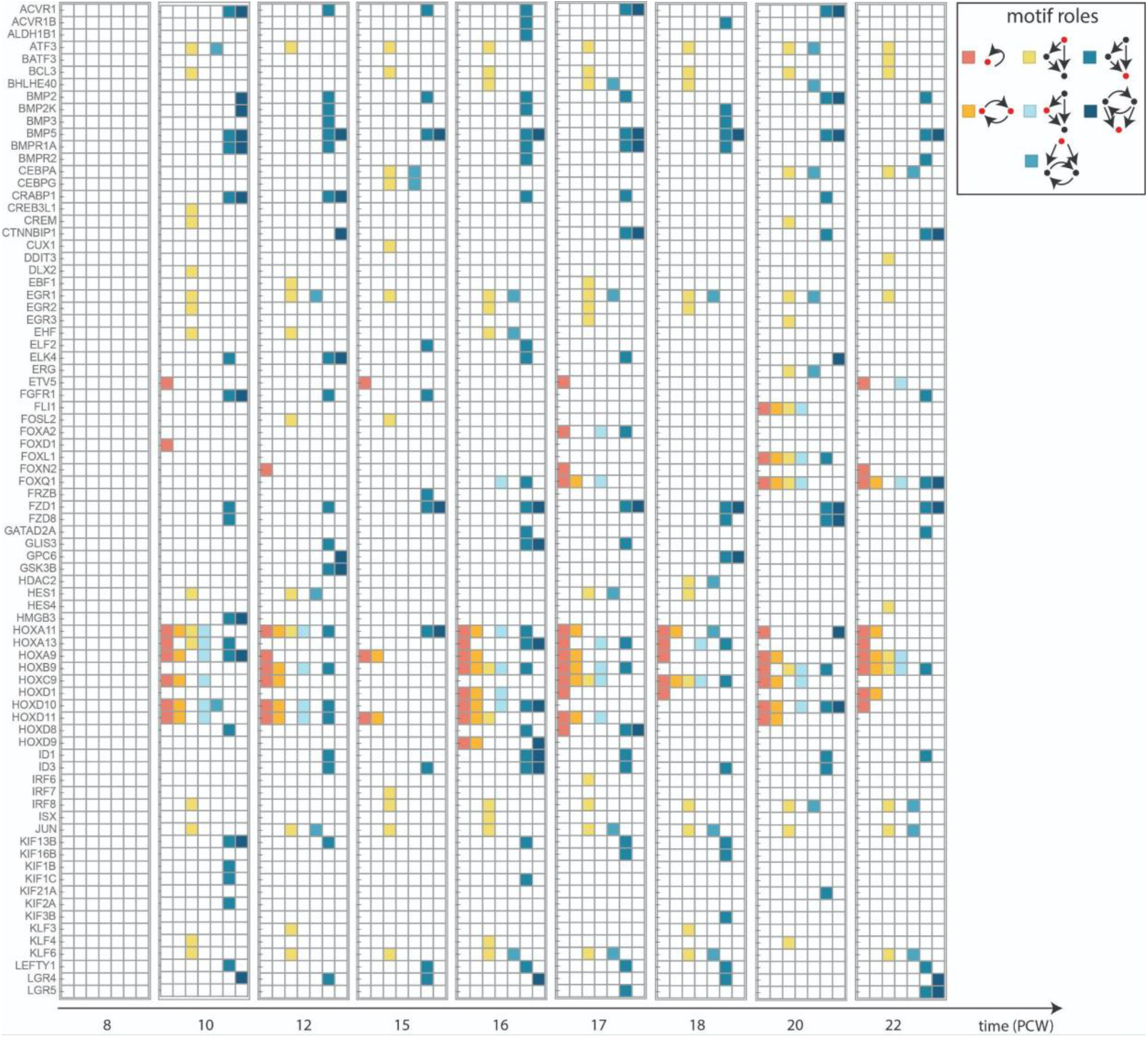
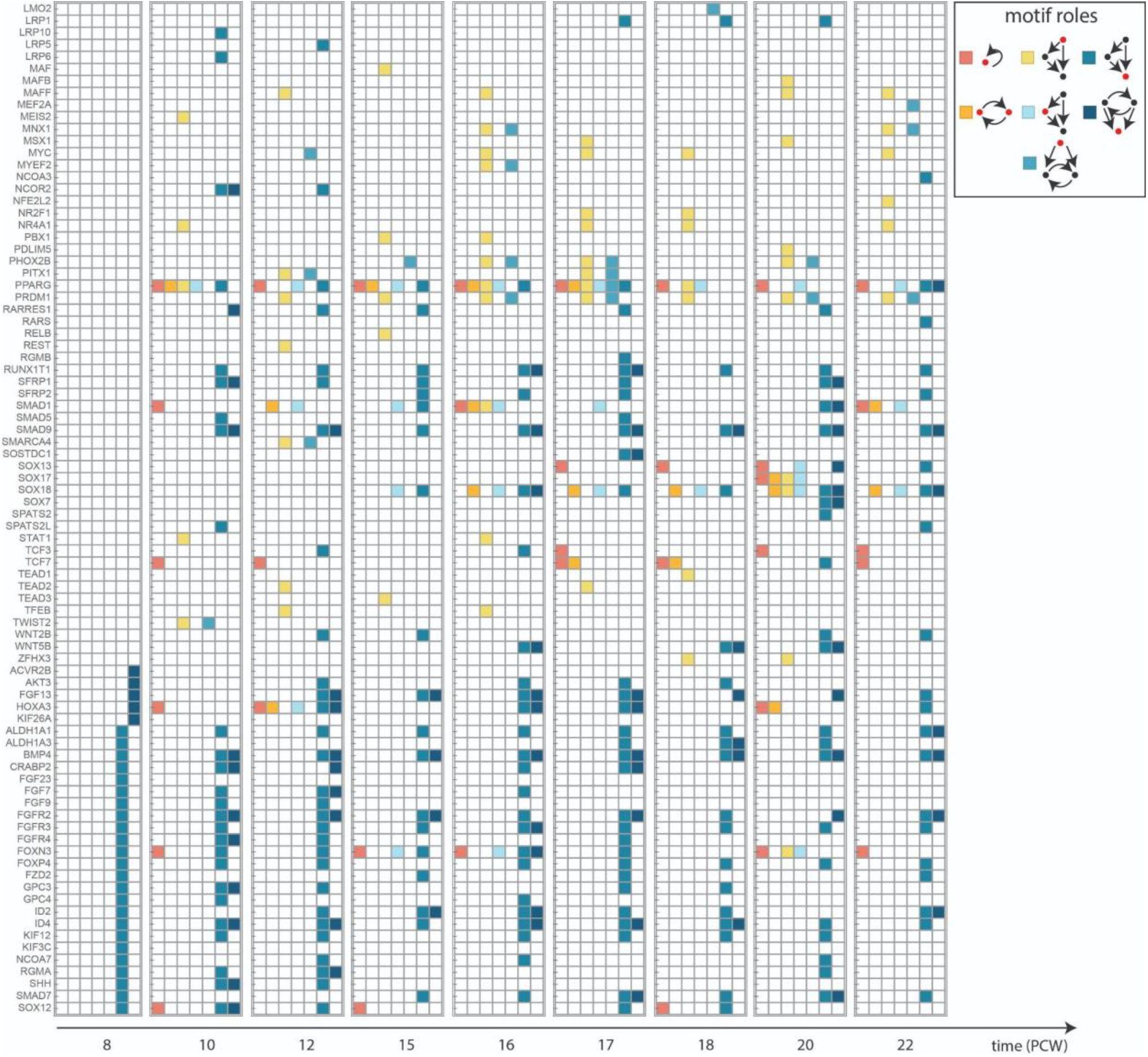
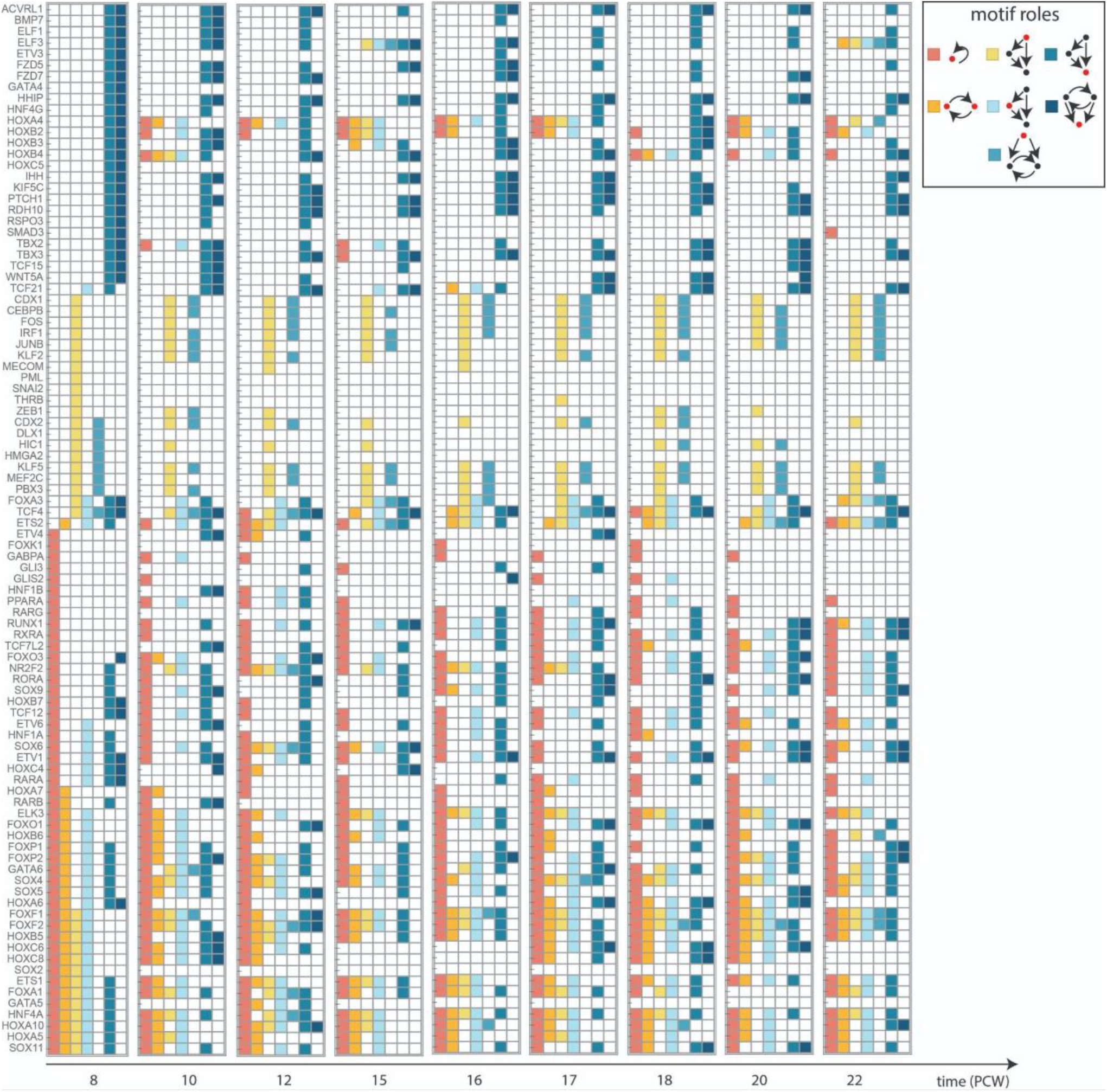
Table of genes participating in network motifs in at least one of the networks inferred from the single-cell data, and their roles within the network motifs in each time point in the data. Every row is a gene, and it is colored by its role within network motifs in each time point.

